## Supplemental material for "Live imaging of excitable axonal microdomains in ankyrin-G-GFP mice"

*for*

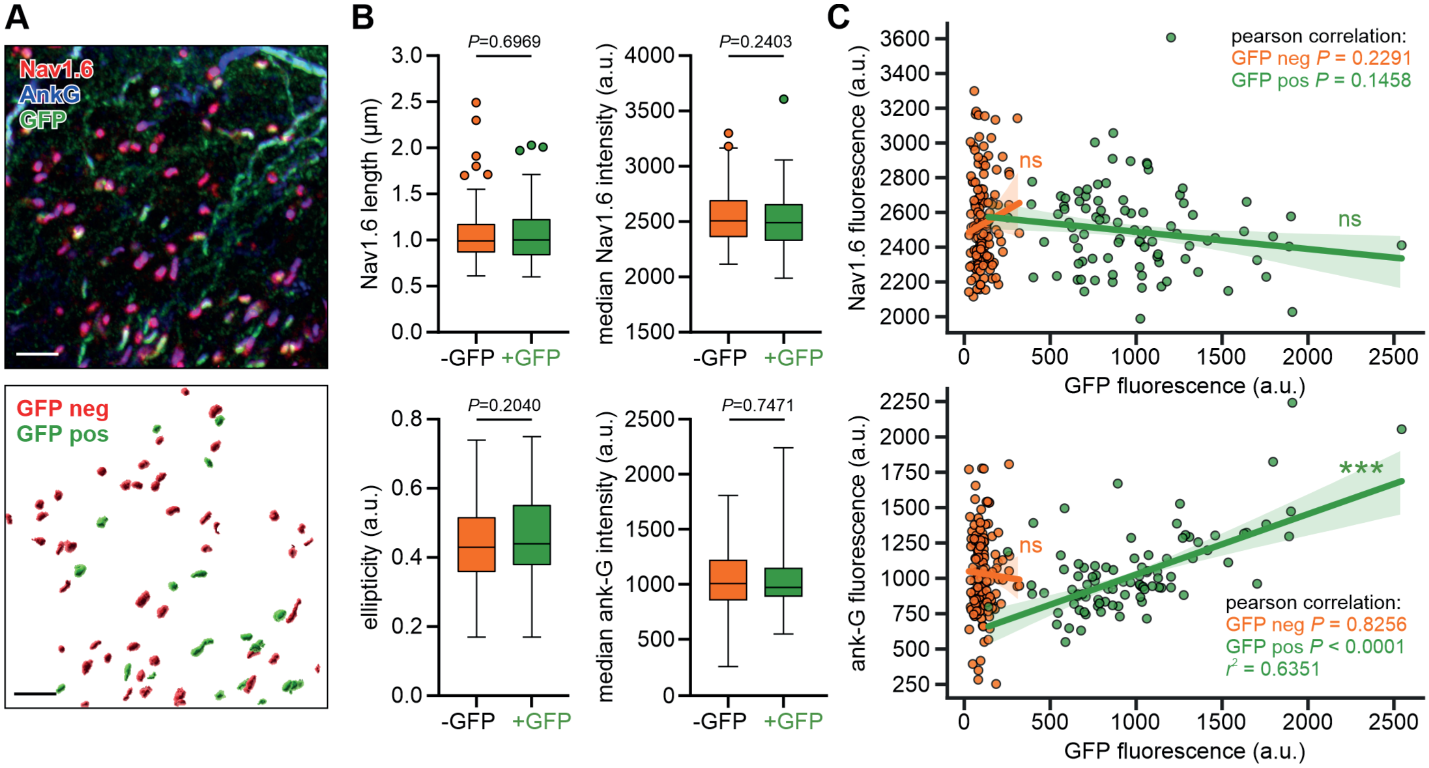

**Supplementary Figure S1: Ank-G-GFP activation and expression in nodes of Ranvier do not alter node morphology**

**A** Top panel shows a cryosection of neocortical white matter from an ankyrin-G-GFP x CaMKIIa-Cre mouse, with $GFP^+$ and $GFP^-$ nodes of Ranvier (noR; see Fig. 3A for details). The bottom panel shows an automated 3D reconstruction of noR using Imaris. Nodes were identified via the ankyrin-G channel, classified by the GFP channel, and properties were analyzed in $Na_v1.6$ and ankyrin-G channels. All steps were automated within the Imaris software. Scale bar = 5 µm. **B** Nodes that are $GFP^+$ and $GFP^-$ show no differences regarding their length, ellipticity, and median fluorescence intensity of ankyrin-G or $Na_v1.6$ signals ($n$ = 141 $GFP^-$ nodes, 91 $GFP^+$ nodes, 1 animal, Mann-Whitney U test). **C** *Top* panel: The fluorescence intensity of the sodium channel $Na_v1.6$ did not correlate with ankyrin-G-GFP fluorescence intensity, indicating unchanged levels of sodium channels. *Bottom* panel: The intensity of GFP fluorescence correlates positively with the levels of all ankyrin-G in $GFP^+$ but not $GFP^-$ nodes ($n$ as in B; Pearson correlation details in the graph). This demonstrates that ankyrin-G-GFP does not change the $Na_v1.6$ channel fluorescence intensity and provides a reliable predictor of ankyrin-G levels, though we do not exclude the possibility that native unlabeled ankyrin-G remains in the nodes to some degree.

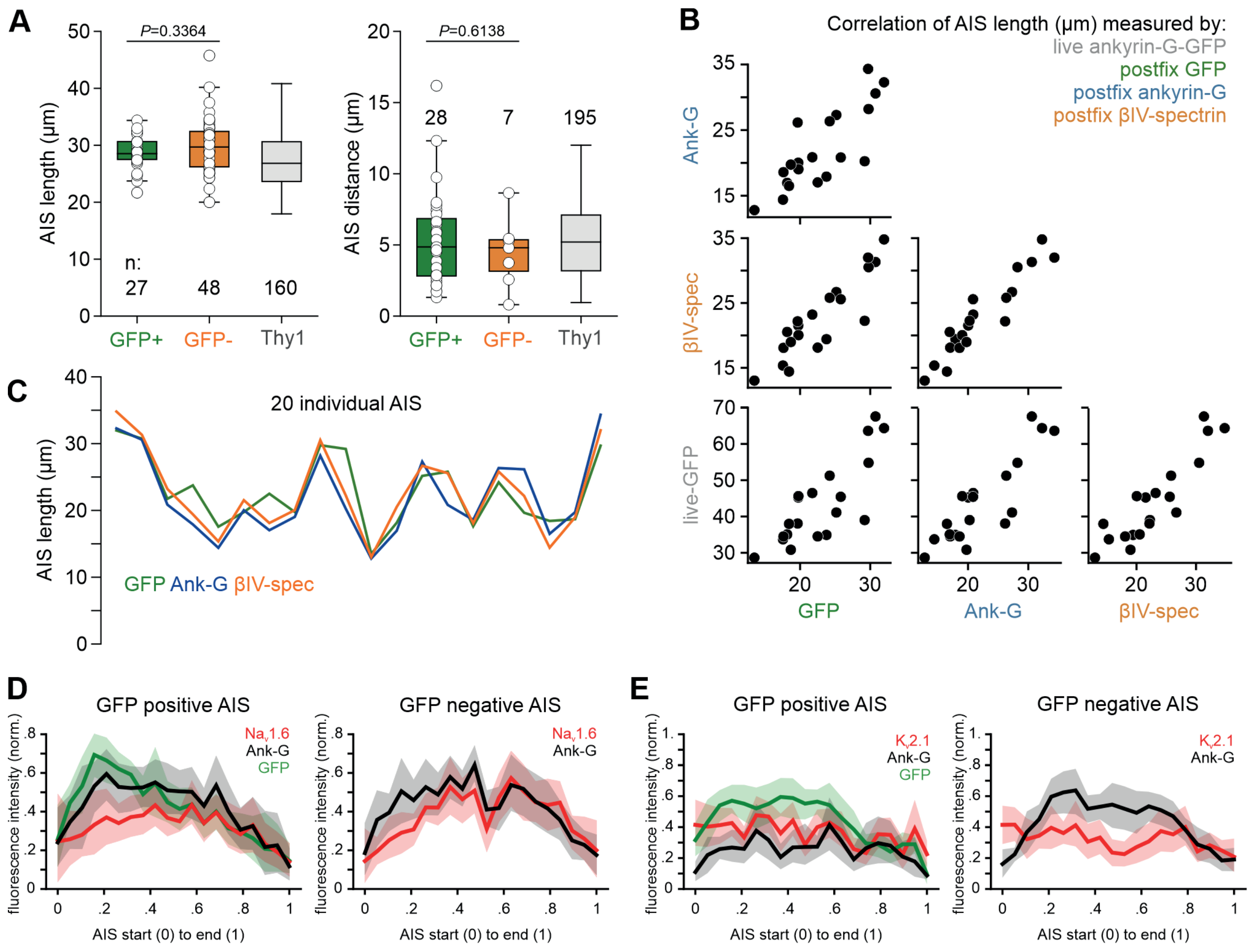

**Supplementary Figure S2: AIS length, position, and molecular composition remains intact after ankyrin-G-GFP expression**

**A** Measurements of length and position of AIS in CA1 pyramidal neurons that are either GFP positive (green) or negative (orange). The inclusion of GFP into the ankyrin-G gene does not significantly change AIS length (*left* panel) or distance to the soma (*right* panel). Sample preparation was conducted as outlined in Fig. 7 and AIS measurements were based on the βIV-spectrin signal ($n$ = 3 animals). An independent Thy1-GFP control line (grey, $n$ = 3 animals) was used as an additional control. The number of individual AIS (white circles) and $P$ values are given within the graph ($t$-test for AIS length; Mann – Whitney U-test for AIS distance). **B** Correlation of AIS length measured live via the ankyrin-G-GFP signal in a patch clamp chamber (grey) and post-fixation using antibodies against GFP, ankyrin-G, and βIV-spectrin in CA1 pyramidal neurons (sample preparation as in Fig. 7, $n$ = 20 AIS, 1 animal). **C** Alternative visualization of data from panel B (post-fixation). Measurements for AIS signals using antibodies against GFP, ankyrin-G, and βIV-spectrin show comparable lengths. **D** Immunosignals from $Na_v$1.6 channels retain their fluorescence intensity across the AIS after the expression of ankyrin-G-GFP. AIS length was normalized from start to end of the ankyrin-G signal and fluorescence intensity from lowest to highest. Tissue preparation as described in Fig. 4A ($n$ = 10 GFP$^+$ and 10 GFP$^-$ AIS). **E** Line plots of $K_v$2.1 and GFP fluorescence intensities along the AIS signal indicate no change in $K_v$2.1 expression. Normalization as in panel D. Tissue preparation as described in Fig. 4B ($n$ = 10 GFP$^+$ and 10 GFP$^-$ AIS).

**Supplementary Table 1** Summary of statistics for passive and active properties (related to Fig. 7)

| stats | wildtype N | mean | median | SD | AnkG-GFP N | mean | median | SD | AnkG-GFP+Cre N | mean | median | SD | ANOVA p(A) / p(KW) | -Cre / +Cre ttest / M-Whit |
|---|---|---|---|---|---|---|---|---|---|---|---|---|---|---|
| RMP | 28 | -63.29 | -63.56 | 3.78 | 22 | -62.57 | -63.43 | 4.62 | 42 | -64.14 | -63.32 | 4.17 | 0.3441 | 0.1733 |
| Rinput | 28 | 268.60 | 251.50 | 95.51 | 23 | 226.10 | 203.70 | 89.55 | 42 | 253.00 | 234.40 | 95.75 | 0.0952 | 0.1634 |
| Ih sag | 28 | 22.25 | 22.01 | 8.67 | 23 | 22.84 | 23.30 | 8.64 | 42 | 25.06 | 28.05 | 9.91 | 0.2529 | 0.3469 |
| rheobase | 28 | 52.16 | 44.86 | 30.90 | 23 | 70.86 | 69.81 | 36.43 | 41 | 69.31 | 69.81 | 37.54 | 0.0654 | 0.9917 |
| ISI (ratio 1/6) | 27 | 2.55 | 2.32 | 1.00 | 22 | 2.39 | 2.11 | 0.68 | 42 | 1.98 | 1.86 | 0.47 | 0.0038 | 0.0123 |
| maxAP | 27 | 12.04 | 11.00 | 2.59 | 22 | 9.77 | 10.00 | 2.05 | 42 | 10.52 | 10.00 | 2.48 | 0.0044 | 0.2278 |
| IhalfmaxAP | 27 | 102.40 | 96.82 | 39.97 | 22 | 122.20 | 120.10 | 32.56 | 42 | 122.90 | 116.30 | 46.33 | 0.0576 | 0.6471 |
| AP thesh | 28 | -39.00 | -38.45 | 2.75 | 23 | -38.30 | -38.70 | 3.75 | 42 | -36.91 | -36.54 | 3.41 | 0.0163 | 0.0457 |
| AP amp. | 28 | 84.98 | 86.78 | 10.90 | 23 | 88.85 | 90.03 | 7.73 | 42 | 90.29 | 91.51 | 6.92 | 0.0396 | 0.4537 |
| AP hw | 28 | 2.05 | 2.04 | 0.26 | 23 | 1.91 | 1.88 | 0.21 | 42 | 1.94 | 1.91 | 0.21 | 0.0546 | 0.5371 |
| AP rise | 28 | 0.54 | 0.54 | 0.12 | 23 | 0.50 | 0.49 | 0.07 | 42 | 0.46 | 0.45 | 0.08 | 0.0013 | 0.0103 |
| AP decay | 28 | 2.19 | 2.19 | 0.41 | 23 | 2.07 | 2.03 | 0.31 | 42 | 2.19 | 2.20 | 0.27 | 0.2392 | 0.1174 |
| EPSP amp. | 27 | 0.59 | 0.59 | 0.08 | 22 | 0.59 | 0.59 | 0.10 | 42 | 0.61 | 0.58 | 0.12 | 0.8246 | 0.5555 |
| EPSP frequ. | 27 | 0.47 | 0.31 | 0.61 | 22 | 0.49 | 0.32 | 0.50 | 42 | 0.40 | 0.25 | 0.41 | 0.6986 | 0.3858 |

multiple comparisons:
significant pairs:

| | |
|---|---|
| ISI (ratio 1/6) | Ank-G-GFP+Cre is different from both others |
| maxAP | wildtype is different from both Ank-G-GFP groups |
| AP thesh | wildtype vs. ankg□GFP+cre |
| AP amp. | wildtype is different to ankG□GFP+Cre |
| AP rise | wildtype vs ankG□GFP+Cre |

**Supplementary Table 2** Summary of Cre viruses and Cre driver lines

| Neuron population | Cre virus / line | Common name | Source, strain |
|---|---|---|---|
| Unspecific | FUW-nGFP::Cre (lentiviral vector), CMV promotor | nGFP-Cre, NA | Gift from the Südhof lab, Stanford University, CA, USA |
| Unspecific | FUW-nGFP:: ΔCre (lentiviral vector), CMV promotor | nGFP-ΔCre, NA | Gift from the Südhof lab, Stanford University, CA, USA |
| Excitatory neurons | pENN.AAV.CamKII 0.4.Cre.SV40 | CaMKII-Cre | Addgene viral prep #105558-AAV5, Lot: v7050 |
| Neurons (unspecific) | ENN.AAV.hSyn.Cre.WPRE.hGH | Synapsin-Cre | Addgene viral prep #105553-AAV1, Lot: v75882 |
| Inhibitory neurons | AAV1-hDlx-Flex-dTomato-Fishell_7 | hDlx-tdTomato | Addgene viral prep #83894-AAV1 |
| Neurons (unspecific) | pAAV-hSyn-Cre-P2A-dTomato | Synapsin-Cre-tdTomato | Addgene viral prep 107738-AAVrg, Lot: v75881 |
| Excitatory neurons | B6.Cg-Tg(Camk2a-Cre)T29-1Stl/J | T29-1<br>IMSR_JAX:005359 | The Jackson Laboratory<br>Strain # 005359 |
| Parvalbumin-pos. interneurons with tdTomato | B6;129P2-Pvalb$^{tm1(Cre)Arbr}$/J<br>crossed with<br>B6.Cg-GT(ROSA)26Sor$^{tm14(CAG-tdTomato)/Hze}$/J | B6 PV$^{Cre}$<br>IMSR_JAX:017320<br><br>Ai14<br>IMSR_JAX:007914 | The Jackson Laboratory<br>Strain # 017320<br><br>Strain # 007914 |
| Unspecific | AAV-retro/2-hSyn1-mCherry-iCre-WRPE-hGHp(A) | Retro-mCherry-Cre | University of Zürich, Viral Vector Facility, v230 |

**Supplementary Table 3** Specification of primary and secondary antibodies (catalog number, working dilution, previously conducted controls, sources, Research Resource Identification Portal (RRID) code and references where available).

| Primary Antibody Clone/type | Dilution | Reported controls | | | | Source & Catalog Number RRID or other reference |
|---|---|---|---|---|---|---|
| | | KO | IF | IP | WB | |
| *Ankyrin-G* (rb) | 1:500 | | X | | X | Santa Cruz Biotechnology, Heidelberg, Germany; sc-28561<br>AB_ 633909 |
| *Ankyrin-G* (gp) | 1:1000 | | X | | X | Synaptic Systems, Göttingen, Germany<br>386 005<br>[1] |
| *Ankyrin-G* (ms)<br>N106/36 | 1:500 | X | X | | X | UC Davis/NIH NeuroMab Facility, CA, USA; 73-146<br>AB_2315803 |
| *ank-G C-terminus* | 1:200 | | X | | | Santa Cruz Biotechnology, Heidelberg, Germany; sc-28561; discontinued |
| *ßIV-spectrin* (rb) | 1:1000 | X | X | | X | Self-made<br>[2-4] |
| *Caspr* (ms)<br>K65/35 | 1:500 | X | X | X | X | UC Davis/NIH NeuroMab Facility, CA, USA; 75-001<br>AB_2083496 |
| *FGF14* (ms)<br>N56/21 | 1:500 | X | X | X | X | UC Davis/NIH NeuroMab Facility, CA, USA; 75-096<br>AB_2104060 |
| *GFP* (ch) | 1:1000 | | X | | | Acris Antibodies GmbH, Hiddenhausen, Germany;<br>AP20142PU-N<br>AB_10756183 |
| *Iba1* (rb) | 1:2000 | | X | | | Wako, Neuss, Germany; 019-19741<br>AB_839504 |
| $K_v$*1.2* (ms)<br>K14/16 | 1:200 | X | X | X | X | UC Davis/NIH NeuroMab Facility, CA, USA; 73-008<br>AB_2296313 |
| $K_v$*2.1* (ms)<br>K89/34 | 1:1000 | X | X | | X | UC Davis/NIH NeuroMab Facility, CA, USA; 75-014-020<br>AB_2877280 |
| *Nav1.6 (rb)* | 1:2000 | | X | | X | Alomone Labs, Jerusalem, Israel<br>ASC-009<br>AB_2040202 |
| *NeuN (ms)*<br>*A60* | 1:500 | | X | | X | Millipore, Temecula, CA, USA; MAB377<br>AB_2314889 |
| *NeuN (gp)* | 1:2000 | | X | | | Synaptic Systems, Göttingen, Germany 266 004<br>AB_2619988 |
| *Parvalbumin (rb)* | 1:500 | | X | | X | Swant Inc., Marly, Switzerland<br>PV27<br>AB_2631173 |
| *Synaptopodin (gp)* | 1:500 | | X | | X | Synaptic Systems, Göttingen, Germany 163 004<br>AB_10549419 |
| *TRIM46 (ms)*<br>*SMP14*<br>*(aka MDM2)* | 1:500 | | X | | X | Santa Cruz Biotechnology, Heidelberg, Germany; sc-965<br>AB_627920 |
| *vGAT (ch)* | 1:1000 | | X | | | Synaptic Systems, Göttingen, Germany 131 006<br>AB_2619820 |

*KO* absence of immunosignal in knock out animals, *IF* immunofluorescence, *IP* immuno-precipitation, *WB* western blot, *rb* rabbit, *ms* mouse, *gp* guinea pig, *ch* chicken

**Supplementary Table 3 (continued)** Specification of primary and secondary antibodies (catalog number, working dilution, previously conducted controls, sources, Research Resource Identification Portal (RRID) code and references where available).

| **Secondary Antibody Clone/type** | **Dilution** | **Reported controls** | | | | **Source & Catalog Number RRID or other reference** |
|---|---|---|---|---|---|---|
| | | **KO** | **IF** | **IP** | **WB** | |
| *FluoTag-X4-anti GFP-StarRED* 1H1/1B2 | 1:500 | | | | | NanoTag Technologies, Göttingen, Germany; N0304 AB_2744631 |
| *gt anti mouse abberior STAR 580* | 1:100 | | | | | Abberior GmbH, Göttingen, Germany; ST580-1001 AB_2923543 |
| *gt anti rabbit abberior STAR 580* | 1:100 | | | | | Abberior GmbH, Göttingen, Germany; ST580-1002 AB_2910107 |
| *gt anti ch Alexa Fluor 488* | 1:1000 | | | | | Molecular Probes, Thermo Fisher, Karlsruhe, Germany; A-11039 AB_ 42924 |
| *gt anti ms Alexa Fluor 568* | 1:1000 | | | | | Molecular Probes, Thermo Fisher, Karlsruhe, Germany; A-11004 AB_143162 |
| *gt anti rb Alexa Fluor 568* | 1:1000 | | | | | Molecular Probes, Thermo Fisher, Karlsruhe, Germany; A-21069 AB_141416 |
| *gt anti gp Alexa Fluor 568* | 1:1000 | | | | | Molecular Probes, Thermo Fisher, Karlsruhe, Germany; A-11075 AB_ 141954 |
| *gt anti ch Alexa Fluor 568* | 1:1000 | | | | | Molecular Probes, Thermo Fisher, Karlsruhe, Germany; A-11041 AB_2534098 |
| *gt anti ms Alexa Fluor 594* | 1:500 | | | | | Molecular Probes, Thermo Fisher, Karlsruhe, Germany; A-21125 AB_141593 |
| *gt anti rb Alexa Fluor 594* | 1:500 | | | | | Molecular Probes, Thermo Fisher, Karlsruhe, Germany; A-11012 AB_141359 |
| *gt anti ms Alexa Fluor 647* | 1:500 | | | | | Molecular Probes, Thermo Fisher, Karlsruhe, Germany; A AB_2535804-21235 |
| *gt anti rb Alexa Fluor 647* | 1:500 | | | | | Molecular Probes, Thermo Fisher, Karlsruhe, Germany; A-21244 AB_2535812 |
| *gt anti gp Alexa Fluor 647* | 1:500 | | | | | Molecular Probes, Thermo Fisher, Karlsruhe, Germany; A-21450 AB_2735091 |
| *gt anti ch Alexa Fluor 647* | 1:500 | | | | | Molecular Probes, Thermo Fisher, Karlsruhe, Germany; A-21449 AB_2535866 |
| *Alexa Streptavidin 568* | 1:1000 | | | | | Molecular Probes, Thermo Fisher, Karlsruhe, Germany; *S11226* AB_2315774 |

*KO* absence of immunosignal in knock out animals, *IF* immunofluorescence, *IP* immuno-precipitation, *WB* western blot, *rb* rabbit, *ms* mouse, *gp* guinea pig, *ch* chicken

**Supplementary Table 4** Summary of fixation and blocking reagents for all immunofluorescence experiments.

| **Sample** | **Fixation**<br>**Time / DIV** | **Post Fix** | **Blocking buffer**<br><br>**Incubation buffer** |
|---|---|---|---|
| Isolated hippocampal neurons | 4% PFA, 15 min<br>DIV 19 | No | **B:** 1% BSA in PBS. Prior to blocking, quench in PBS with 100 mM glycine, 100 mM ammonium chloride (5 min) and application of 0.1% Triton X-100 for 5 min<br><br>**I:** PBS |
| Hippocampal OTC | 4% PFA, 30 min | No | **B:** 0.1% Triton X-100, 1% BSA, 0.2% fish skin gelatine, in 1 x PBS<br><br>**I:** same |
| *Ex vivo* acute slices | 2% PFA, 90 min | No | **B:** 0.3% Triton X-100, 5% normal goat serum in 1 x PBS<br><br>**I:** 0.2% Triton X-100, 1% normal goat serum in 1 x PBS |
| Whole mount retina | 4% PFA, 15 min<br>>P55 | No | **B:** 0.5% Triton X-100, 0.2% BSA, 0.02% sodium azide in 1x PBS<br><br>**I:** 1% Triton X-100, 10% FCS and 0.02% sodium azide in 1x PBS |
| Cryosections | Perfusion, 15 min<br>2% PFA (for ion channels)<br>4% PFA (for all others)<br>>P28 | No | **B:** 1% BSA, 0.2% fish skin gelatine, 0.1% Triton X-100 in 1 x PBS<br><br>**I:** same |

*PFA* paraformaldehyde, *min* minutes, *DIV* days in vitro, *BSA* bovine serum albumin, *PBS* phosphate buffered saline, *FCS* fetal calf serum
